# Targeting the endolysosomal host-SARS-CoV-2 interface by clinically licensed functional inhibitors of acid sphingomyelinase (FIASMA) including the antidepressant fluoxetine

**DOI:** 10.1101/2020.07.27.222836

**Authors:** Sebastian Schloer, Linda Brunotte, Jonas Goretzko, Angeles Mecate-Zambrano, Nadia Korthals, Volker Gerke, Stephan Ludwig, Ursula Rescher

**Affiliations:** Institute of Medical Biochemistry, Center for Molecular Biology of Inflammation, and “Cells in Motion” Interfaculty Centre, University of Muenster, Von-Esmarch-Str. 56, D-48149, Muenster, Germany; Institute of Virology, Center for Molecular Biology of Inflammation, and “Cells in Motion” Interfaculty Centre University of Muenster, Von-Esmarch-Street 56, D-48149 Muenster, Germany; Interdisciplinary Centre for Clinical Research, University of Muenster, D-48149 Muenster, Germany

**Keywords:** SARS-CoV-2, IAV, FIASMA, fluoxetine, viral entry, endolysosomal interference

## Abstract

The Corona Virus Disease 2019 (COVID-19) pandemic caused by the Severe Acute Respiratory Syndrome Related Coronavirus 2 (SARS-CoV-2) is a global health emergency. As only very limited therapeutic options are clinically available, there is an urgent need for the rapid development of safe, effective, and globally available pharmaceuticals that inhibit SARS-CoV-2 entry and ameliorate COVID-19. In this study, we explored the use of small compounds acting on the homeostasis of the endolysosomal host-pathogen interface, to fight SARS-CoV-2 infection. We find that fluoxetine, a widely used antidepressant and a functional inhibitor of acid sphingomyelinase (FIASMA), efficiently inhibited the entry and propagation of SARS-CoV-2 in the cell culture model without cytotoxic effects and also exerted potent antiviral activity against two currently circulating influenza A virus subtypes, an effect which was also observed upon treatment with the FIASMAs amiodarone and imipramine. Mechanistically, fluoxetine induced both impaired endolysosomal acidification and the accumulation of cholesterol within the endosomes. As the FIASMA group consists of a large number of small compounds that are well-tolerated and widely used for a broad range of clinical applications, exploring these licensed pharmaceuticals may offer a variety of promising antivirals for host-directed therapy to counteract enveloped viruses, including SARS-CoV-2 and COVID 19.

## Introduction

The current outbreak of the Severe Acute Respiratory Syndrome Related Coronavirus 2 (SARS-CoV-2) and the resulting Corona Virus Disease 2019 (COVID-19) pandemic clearly show the potential global threat of a newly emerging pathogen. The worldwide spread and the immense number of infected people worldwide pose not only a severe risk for the clinical management of individual patients but increasingly for the healthcare systems and the global economy in general ^1^. While coronaviruses typically circulate in the human population and cause mild respiratory illness ^2^, COVID-19 elicited by the zoonotic SARS-CoV-2 which is also transmitted from person to person ^3^ is a much more severe disease, and elderly patients and those with chronic medical conditions are at higher risk of serious illness ^4–7^. As there are only very limited treatment options so far, unprecedented global efforts are made to limit transmission and develop tools and therapeutic strategies to combat infection and improve the outcome of the disease. Besides the development of efficient vaccines, therapeutic antibodies, and virus-targeting drugs, the identification of host cell factors that are required for the successful SARS-CoV-2 replication process might offer effective drug targets. Ideally, existing licensed drugs would be repurposed as a faster and cheaper pathway to develop new SARS-CoV-2 antivirals.

Typically for coronaviruses, the SARS-CoV-2 positive-strand RNA genome is packaged within the enveloped capsid^8^. Entry into host cells is induced by binding of the viral surface spike (S) protein, which protrudes from the virus envelope to the angiotensin-converting enzyme 2 (ACE2) on the surface of the host cell ^9^. According to its classification as a class I viral fusion protein, proteolytic cleavage is required to release the S protein fusogenic peptide, thus inducing the actual fusion event with the host cell membrane and the transfer of the viral genome into the cytosol ^9^. S protein can be primed by several cellular proteases. Dependent on protease availability, cleavage can occur via the transmembrane protease serine 2 (TMPRSS2), leading to fusion with the plasma membrane, and virus particles that are endocytosed following ACE2 binding can use endosome-residing proteases for fusion with the endosomal membrane. Both pathways are shown to contribute to the SARS-CoV-2 infection process ^9^.

For enveloped viruses that use the endocytic pathway in the entry process, the endosome is an important host-pathogen interface and the conditions of the endosomal environment critically affect the virus-endosome fusion and thus, endosomal escape ^10,11^. Our previous studies identified the host cell cholesterol homeostasis as a crucial component of the infection process of endocytosed enveloped viruses. Importantly, we verified that the increase in late endosomal cholesterol is embedded in the interferon response against incoming influenza A virus (IAV) particles ^12^, that this barrier has antiviral capacities when raised pharmacologically ^13,14^, and that this druggable host cell target can be exploited through repurposed drugs in a preclinical murine infection model ^13^.

Endosomal lipid balance is controlled via a broad range of feedforward as well as feedback circuits and is part of the intricate lipid sorting and degradation system that operates in late endosomes/lysosomes (LELs) ^15^. Importantly, structurally unrelated small cationic amphiphilic molecules, such as the sterol derivative U18666A and a broad range of commonly used and well-tolerated human medications including the clinically licensed anti-depressant fluoxetine, are known to induce accumulation of cholesterol in LELs ^16^. Whereas U18666A has been identified as a direct inhibitor of the endolysosomal cholesterol transport protein NPC1 ^17^, the mode of action of other members of this drug class is less clear and might involve blockade of the sphingomyelin converting acid sphingomyelinase (ASMase) within the LELs ^16^. In this study, we explored the impact of disturbed ASMase activity and endolysosomal cholesterol accumulation on SARS-CoV-2 infection in Vero E6 cells commonly used to investigate SARS-CoV-2 infection, and in polarized bronchial Calu-3 cell monolayers, a lung cell model for productive SARS-CoV-2 infection. Based on our previous results on the antiviral capacity of dysregulated endolysosomal cholesterol contents, we reasoned that clinically licensed drugs associated with lipid storage in LELs might be exploited as repurposed antivirals for host-directed therapy against SARS-CoV-2. Based on this rationale, we assessed the antiviral potential of treatment with fluoxetine, a selective serotonin reuptake inhibitor (also known under the trade-name Prozac) commonly used to treat major depression and related disorders ^18^ on IAV and SARS-CoV-2 infection. Our results show that fluoxetine pretreatment inhibited IAV infection. Most importantly, fluoxetine treatment was able to inhibit acute SARS-CoV-2 infection in a dose-dependent manner up to 99%. Notably, the inhibitory effects were also seen with two other FIASMAs, amiodarone and imipramine. Our data support the hypothesis that the endolysosomal lipid balance is a promising druggable target at the host-virus interface for a wide variety of enveloped viruses, including IAV and SARS-CoV-2. Tapping this vast pool of potential antivirals for drug repurposing might be a fast and powerful approach to fight a broad range of pathogens with functionally similar modes of action, which is especially important for newly emerging viruses such as SARS-CoV-2.

## Material and Methods

### Cells and treatment

The human bronchioepithelial cell line Calu-3, the Madin-Darby canine kidney (MDCK) II cells, and the Vero E6 cell line were cultivated in Dulbecco’s modified Eagle’s medium (DMEM) supplemented with 10% standardized fetal bovine serum (FBS Advance; Capricorne), 2 mM L-glutamine, 100 U/mL penicillin, and 0.1 mg/mL streptomycin. All cell lines were cultured in a humidified incubator at 37°C and 5% CO_2_. Calu-3 were polarized on semipermeable membrane supports as described ^14^. Fluoxetin (5 mM, Sigma) and the NPC1 small molecule inhibitor U18666A (10 mg/mL, Biomol) and amiodarone (5mM, Sigma) were solubilized in DMSO and imipramine was solubilized in water (100mM, Sigma). Cells were treated with 2 μg/mL and /10 µg/mL of U18666A or the indicated fluoxetine, amiodarone and imipramine concentration either for 16 hours before infection or one hour post-infection (p.i) at the indicated drug concentrations for the entire 48 hours infection period.

### Cytotoxicity assay

Calu-3 and Vero cells were treated at the indicated concentrations with the solvent DMSO, U18666A, fluoxetine, amiodarone and imipramine. The strong cytotoxic agent staurosporine (1 μM) served as a positive control for cytotoxic effects. Following 48 hours of treatment, MTT 3-(4,5-dimethylthiazol-2-yl)-2,5-diphenyltetrazolium bromide (Sigma) was added to the cells for 4 h. The supernatant was aspirated, DMSO was added to the cells for 5 min, and subsequently, the OD_562_ was measured ^34^.

### Filipin staining and colocalization analysis

Cells were fixed with 4% PFA in PBS at room temperature for 10 min, subsequently washed twice with PBS and blocked with 2% BSA in PBS for 60 min. To visualize free cellular cholesterol, cells were stained with filipin (complex from Streptomyces filipinensis, 1.25 mg/mL, Sigma) for 2 hours. Late endosomal/lysosomal compartments were identified by staining the endolysosomal marker protein CD63 with the mouse monoclonal anti-CD63 antibody H5C6 (1:300 in 2% BSA in PBS) for 90 min. H5C6 was deposited to the DSHB by August, J.T. / Hildreth, J.E.K. (DSHB Hybridoma Product H5C6). The secondary AlexaFluor594-coupled anti-mouse antibody was from Thermo Fisher Limited. Confocal microscopy was performed with an LSM 780 microscope (CarlZeiss, Jena, Germany) equipped with a Plan-Apochro-mat 63x/1.4 oil immersion objective. The colocalization of filipin and CD63 signals was analyzed using the JACoP plugin ^35^ for Fiji ^36^.

### Endosomal pH measurement

Ratiometric fluorescence microscopy and calculation of endosomal pH were done as described ^12^. In brief, cells were exposed to Oregon green 488 (OG488)-labeled and tetramethylrhodamine (TMR)-labeled dextran (Invitrogen) (10 kDa) for 60 min, followed by a 60 min chase. Cells were then washed and kept in HEPES-buffered Hanks’ balanced salt solution (HBSS) at 37°C during image acquisition. The individual epifluorescence signals of each dye were acquired at intervals of 60 s, and the endosomal pH values of the cells were calculated according to the calibration curve generated by applying standard solutions ranging from pH 4.5 to 6.0 to cells.

### Virus infection

Infection with the IAV isolates A/Hamburg/04/2009 (H1N1)pdm09 and A/Panama/2007/99 (H3N2) from the strain collection of the Institute of Virology, University of Münster, Germany, was carried out as previously described ^14^. The SARS-CoV-2 isolate hCoV-19/Germany/FI1103201/2020 (EPI-ISL_463008, mutation D614G in spike protein) was used for infection of Calu-3 cells and Vero E6 cells in infection-PBS (containing 0.2% BSA, 1% CaCl_2_, 1% MgCl_2_, 100 U/mL penicillin and 0.1 mg/mL streptomycin) at MOI 0.1 for 1 hour.

### Plaque assay

To quantify virus production, supernatants of infected cells were collected at the indicated times p.i. and the number of infectious particles was determined by a standard plaque assay. In brief, MDCK cells (for IAV infection) or Vero E6 cells (for SARS-CoV-2 infection) grown to a monolayer in six-well dishes were washed with PBS and infected with serial dilutions of the respective supernatants in infection-PBS for 1 hour at 37°C. The inoculum was replaced with 2x MEM (MEM containing 0.2% BSA, 2 mM L-glutamine 1 M HEPES, pH 7.2, 7.5% NaHCO_3_, 100 U/mL penicillin, 0.1 mg/mL streptomycin, and 0.4% Oxoid agar) and incubated at 37°C. Virus plaques were visualized by staining with neutral red, and virus titers were calculated as plaque-forming units (PFU) per mL.

### Single-cycle infection assay

Calu-3 cells were infected with SARS-CoV2 and fixed with 4% paraformaldehyde (PFA) in PBS for 15 min at room temperature at the indicated times p.i. To visualize the viral nucleoprotein (NP), cells were permeabilized with 0.2% Triton X-100 in PBS for 30 min, blocked with 2% BSA in PBS for > 1 h, and stained with anti-Nucleocapsid (clone #019 Sino Biological, 1:500 in 2% BSA in PBS) for 2 h, followed by staining with anti-rabbit Alexa Fluor 488 (1:500 in 2% BSA in PBS) for 1 h. Cell nuclei were stained with DAPI (Thermo Scientific, 1:1,000 in 2% BSA in PBS) for 10 min. Cells were analyzed by confocal microscopy using an LSM 800 microscope (Carl Zeiss, Jena, Germany), equipped with a Plan-Apochromat 63x/ 1.4 oil immersion objective.

### Data analysis

A priori power analysis (G*Power 3.1 ^37^) was used to estimate the required sample sizes. Data were analyzed with Prism 8.00 (Graph-Pad). For dose-response curves, virus titers were normalized to percentages of the titers detected in control cells, and drug concentrations were log-transformed. EC values were calculated from the sigmoidal curve fits using a four-parameter logistic (4PL) model. For statistical analysis, significant differences were evaluated using one-way ANOVA followed by Dunnett’s multiple comparison test. **p<0.01, ***p< 0.001, ****p ≤ 0.0001.

## Results

### Fluoxetine treatment impairs SARS-CoV-2 infection in Calu-3 and Vero cells

Building on our previous work on the LEL cholesterol balance as a promising therapeutic target for treatment of enveloped viruses ^12–14,19^, we first assessed the antiviral capability of fluoxetine on IAV infection. Because H1N1 and H3N2 subtypes are currently circulating in the human population, Calu-3 cells pretreated with fluoxetine for 16 h were infected with 0.01 MOI of either A/Hamburg/04/2009 (H1N1)pdm09 or A/Panama/2007/99 (H3N2), and the resulting virus titers were measured 24 h p.i. by a standard plaque assay (Fig. 1A). Fitting of the dose-response curves using nonlinear regression and a four-parameter logistic model revealed for both virus subtypes 50% effective fluoxetine concentrations (EC_50_ values) at approximately 1 µM, and EC_90_ values of approximately 5 - 6 µM.

**Figure 1.**
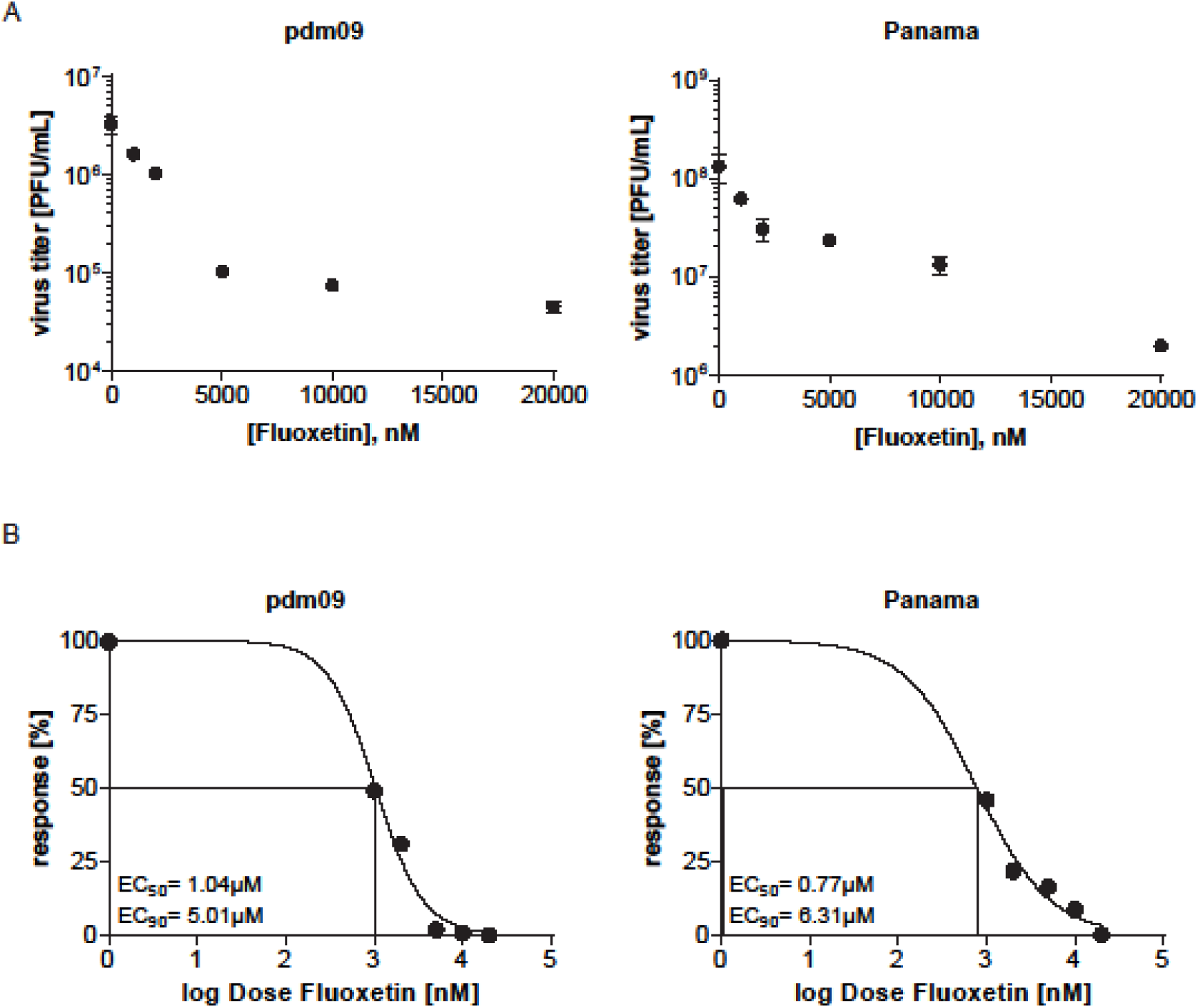
Antiviral potential of fluoxetine treatment against IAV subtypes pdm09 and Panama in Calu-3 cells. (A) Virus titers determined in Calu-3 cells infected with the respective IAV subtype at 0.01 MOI for 24 h. Cells were pretreated with solvent or fluoxetine for 16 h. Data points present mean virus titers ± SEM of three independent experiments. (B) Released viral titers normalized to the control condition and log-transformed fluoxetine concentrations were used to generate the dose-response curves. EC_50_ and EC_90_ values were determined using the 4PL nonlinear regression model.

These results prompted us to assess the antiviral potential of fluoxetine treatment in SARS-CoV-2 infected Vero and Calu-3 cells, two cell lines that are known to produce infectious virus progeny upon SARS-CoV-2 exposure. To analyze the usability of fluoxetine in a more clinically relevant scenario, i.e. an acute infection, we treated the cells 1 h p.i. with a range of fluoxetine concentrations and measured the resulting virus titers 48 h p.i by a standard plaque assay (Fig. 2A). While both Vero and Calu-3 control cells treated with the solvent yielded high viral titers which are in line with previously published results ^9^, dose-dependent inhibition of virus release was observed in both fluoxetine-treated cell lines (Fig. 2A). For both cell lines, 50% effective fluoxetine concentrations (EC_50_ values) were below 1 µM. The 90% inhibition was achieved at ∼ 2 µM for Vero and 4 µM for Calu-3 cells, and higher doses could reduce SARS-CoV-2 titers up to 99% in both cell lines (Fig. 2B).

**Figure 2.**
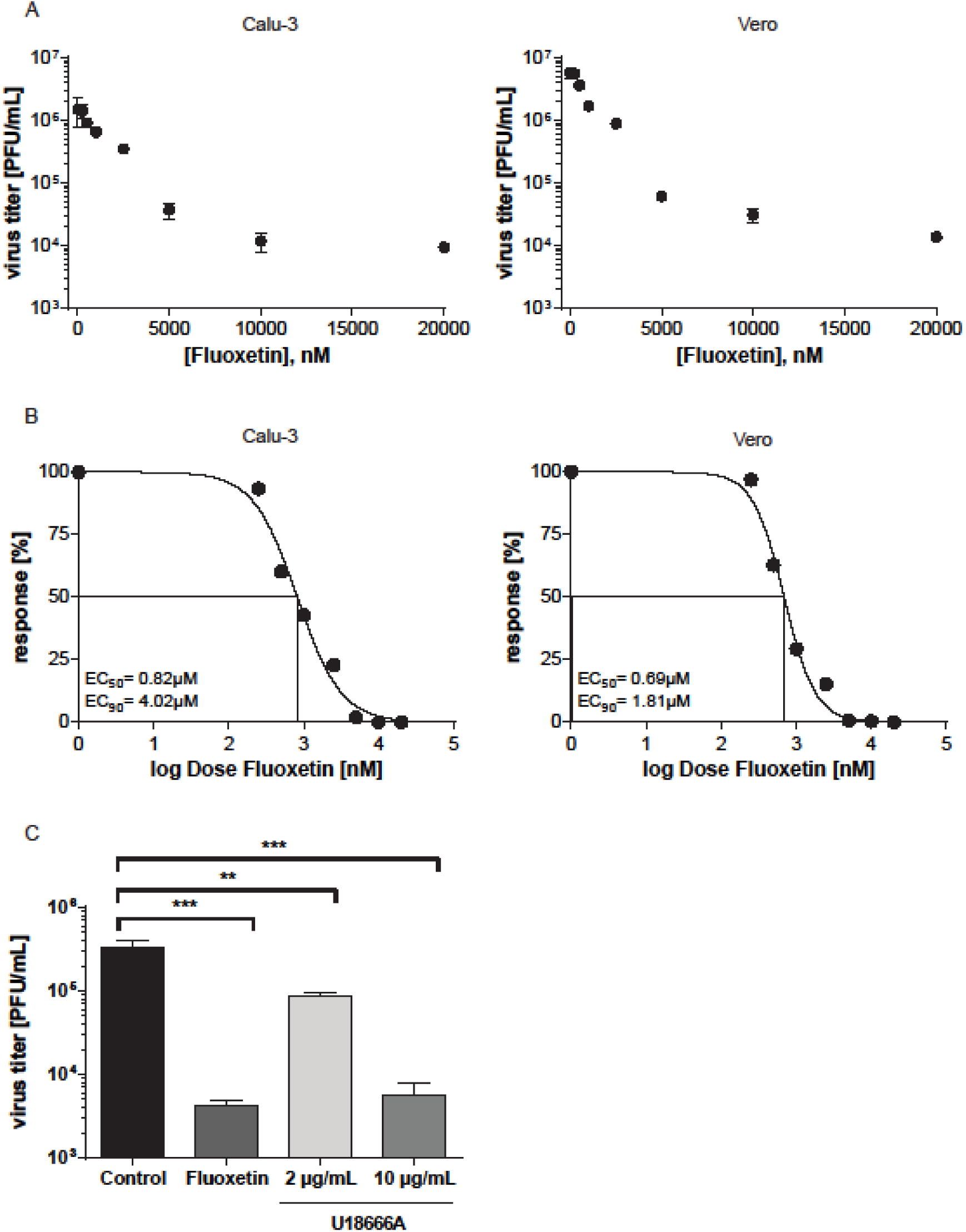
Analysis of anti-SARS-CoV-2 activities of fluoxetine and U18666A treatment in Vero E6 cells and Calu-3 cells. (A) Virus titers determined in Calu-3 and Vero cells infected with SARS-CoV-2 at 0.1 MOI for 48 h. Treatment of infected cells with solvent or fluoxetine was started 1 h p.i. Data points present mean virus titers ± SEM of three independent experiments. (B) To generate the dose-response curves, virus release was normalized to the control condition, fluoxetine concentrations were log-transformed, and nonlinear regression and a 4PL model was used to fit the curves and to determine the EC_50_ and EC_90_ values. (C) Polarized Calu-3 cells grown on semipermeable supports were infected with SARS-CoV-2 isolate at 0.1 MOI for 48 h. Cells were treated 1 h p.i. with 20 µM fluoxetine, and 2 or 10 µg/mL U18666A. Bar graphs represent the mean viral titers ± SEM of three independent experiments. One-way ANOVA followed by by Dunnett’s multiple comparison test. **p ≤ 0.01, ***p ≤ 0.001.

Next, we employed Calu-3 cells polarized on semipermeable membrane support to confirm the results in cells with apicobasal polarity, and thus in a more physiological setting ^14^. Because our previously published results had revealed that increased endolysosomal cholesterol levels act as an effective antiviral barrier for enveloped viruses, including influenza viruses ^12–14,19^, we also tested the impact of a post-infection treatment with U18666A, a small molecule inhibitor of the endolysosomal cholesterol transporter NPC1 ^17^ on SARS-CoV-2 infection. As shown in Fig. 2C, the fluoxetine-mediated inhibition on SARS-CoV2 propagation was faithfully reproduced in the polarized cell model. Of note, U18666A treatment was also antiviral, with a lower concentration of 2 µg/mL reducing the viral titers to 75%, and the higher 10 µg/mL dose displaying a more pronounced antiviral effect of a 99% reduction. Because unwanted cytotoxic effects are one of the main causes for the withdrawal of approved drugs, we measured the impact of U18666A and fluoxetine treatments on Calu-3 and Vero cell viability. As shown in Suppl. Fig. 1, both drug treatments were well tolerated, and cell viability was unaffected over the whole range of the applied dosages and the 48 h treatment period. To evaluate the antiviral potential of the FIASMA group as such, we additionally assessed the antiviral capacity of amiodarone and imipramine. As shown in Fig. 3, both drug treatments reduced SARS-CoV-2 and IAV titers ≥ 90% without any cytotoxic effects, thus supporting our hypothesis.

**Figure 3.**
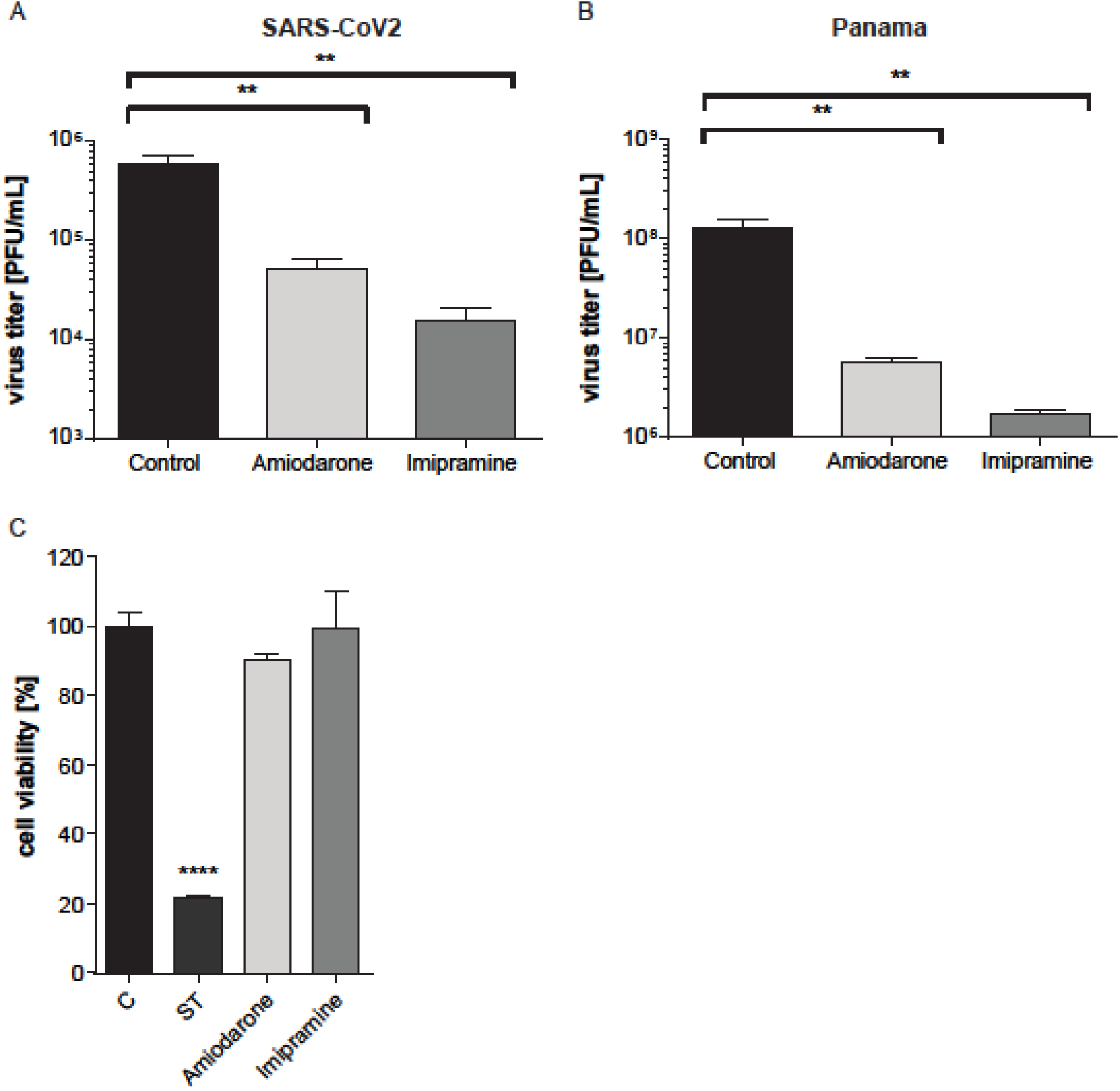
Amiodarone and imipramine as two classic representative of the FIASMA group reduced SARS-CoV2 and IAV Panama titer. Virus titers determined in Calu-3 cells infected with (A) SARS-CoV-2 at 0.1 MOI for 48 h or (B) with the IAV strain Panama at 0.01 MOI for 24 h. Treatment of infected cells with solvent or amiodarone (5 µM) or imipramine (50 µM) was started 1 h p.i. Data points present mean virus titers ± SEM of three independent experiments. (C) Analysis of cell viability. MTT assay of Calu-3 cells treated with the solvent DMSO (C), amiodarone (5 µM) or imipramine (50 µM) for 48 h. The protein kinase inhibitor staurosporine (ST), a strong inducer of cytotoxicity, served as a positive control. Bar graphs represent the mean viral titers ± SEM of three independent experiments. One-way ANOVA followed by Dunnett’s multiple comparison test; **p ≤ 0.01, ****p ≤ 0.0001.

### Fluoxetine treatment is associated with Increased endolysosomal cholesterol storage and dysregulated acidification

Because the pronounced decrease in SARS-CoV-2 titers observed upon treatment with U18666A hinted at an antiviral capacity of dysregulated endolysosomal cholesterol contents in SARS-CoV-2 infection, we next determined whether fluoxetine treatment of Vero and Calu-3 cells was associated with increased endolysosomal cholesterol pools. We acquired z-stacks from confocal microscopy using CD63 as the specific endolysosomal marker protein ^20^ and the polyene macrolide filipin as the cholesterol probe ^12,20^. Filipin is highly fluorescent, binds to unesterified cholesterol in cellular membranes, and the resulting complexes visualize cholesterol by microscopy ^21^. As shown in Fig. 4, for both solvent-treated Calu-3 and Vero cells, filipin stain was diffuse, and only a weak signal was detected in LELs. As expected, blocking endolysosomal cholesterol efflux through inhibition of the major endolysosomal cholesterol transport protein NPC1 with U18666A resulted in the prominent appearance of LELs highly enriched for cholesterol. Of note, fluoxetine at 20 µM also induced the appearance of strong filipin signals in perinuclear vesicles, indicating endolysosomal cholesterol accumulation. For better visualization, the filipin pixels were also color-encoded according to their intensities, resulting in the filipin heatmap (Fig. 4A). For unbiased quantitative image analysis, we calculated the well-established Manders’ coefficient for the co-occurrence of filipin with CD63 signals over the entire volume of the cell. The resulting ratio indicates the total amount of overlapping fluorophore signals and, therefore, indicates changes in endolysosomal cholesterol contents ^12,13^. In line with the images, this quantitative colocalization analysis (Fig 4B), revealed that fluoxetine at a low concentration of 5 µM caused slightly, yet not significantly elevated cholesterol levels in CD63-positive LELs, whereas treatment with 20 µM fluoxetine induced a prominent and significant endolysosomal cholesterol accumulation. The strongest increase in cholesterol levels was observed when cholesterol efflux was directly blocked by the NPC1 inhibitor U18666A.

**Figure 4.**
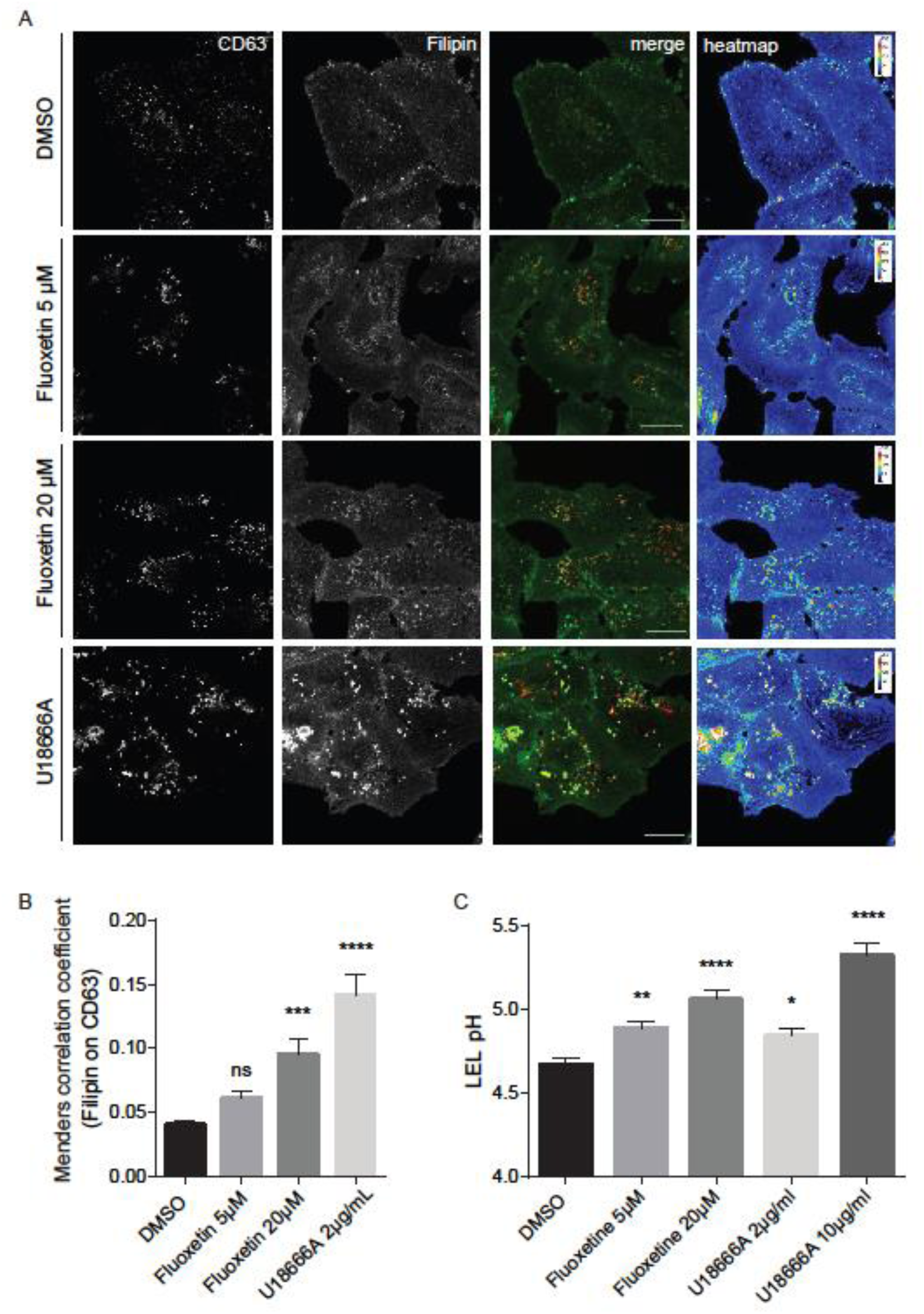
Increased endolysosomal cholesterol storage and dysregulated acidification upon fluoxetine treatment. Vero cells were treated for 16 h with either the solvent DMSO, fluoxetine, or U18666A at the indicated concentrations. (A) Representative 2D maximum intensity projections of entire z-stacks obtained by confocal imaging. LELs were identified via immunodetection of the LEL marker protein CD63 and unesterified cholesterol was visualized using filipin. Representative pseudocolored digital images are shown. To generate the heat maps, filipin-positive pixels were color-encoded according to their intensity values. Scale bar, 20 µm. (B) For each cell, the colocalization of filipin with CD63 quantitated across the entire z-stack was calculated as Manders’ coefficient. Bar graphs represent means ± SEM of 3 individual cells per condition from three independent experiments. (C) Endolysosomal pH values in Calu-3 cells were measured by ratio imaging. Bar graphs present mean pH values ± SEM of 56 cells for each condition from three independent experiments. One-way ANOVA followed by Dunnett’s multiple comparison test; ns, not significant, **p ≤ 0.01, ***p ≤ 0.001, ****p ≤ 0.0001.

A key feature of the late endosomal compartment is the acidic pH. We, therefore, assessed whether fluoxetine treatment impacted endolysosomal pH values in Calu-3 cells, utilizing a quantitative ratiometric fluorescence microscopy assay ^12,13^. In control cells, the measured pH was at a value of 4.7, whereas cells exposed to 10 µg/mL U18666A displayed a significantly reduced acidification, well in agreement with our previously reported data of the impact of higher U18666A concentrations on pH values in LELs ^19^. Of note, the endolysosomal acidification was also affected upon fluoxetine treatment, and this was already observed in cells treated with the low 5 µM concentration (Figure 4C).

To analyze the time point of inhibition, we performed a single-cycle infection assay. Fluoxetine and U18666A were added to the cells 16 h before infection, and cells were infected at MOI 1. Infected cells were then identified 8 hrs p.i. through detection of the SARS-CoV-2 nucleoprotein by immunofluorescence staining. Both drugs impressively reduced the number of NP-positive cells in both Vero and Calu-3 cells (Fig. 5), indicating that their antiviral activities occurred before the release of virus progeny. Detectable NP signals were seen in 17% of the Vero cells. Of note, even a low 5 µM fluoxetine treatment could reduce the number of visibly infected Vero cells to 5%, and the higher dose of 20 µM reduced the levels of infected cells to 2%, indicating a 90% inhibition of cells with detectable SARS-CoV-2 infection. In contrast to the dose-response assays, this assay does not discriminate between different signal strengths, and EC_50_ values are, therefore, not directly comparable.

**Figure 5.**
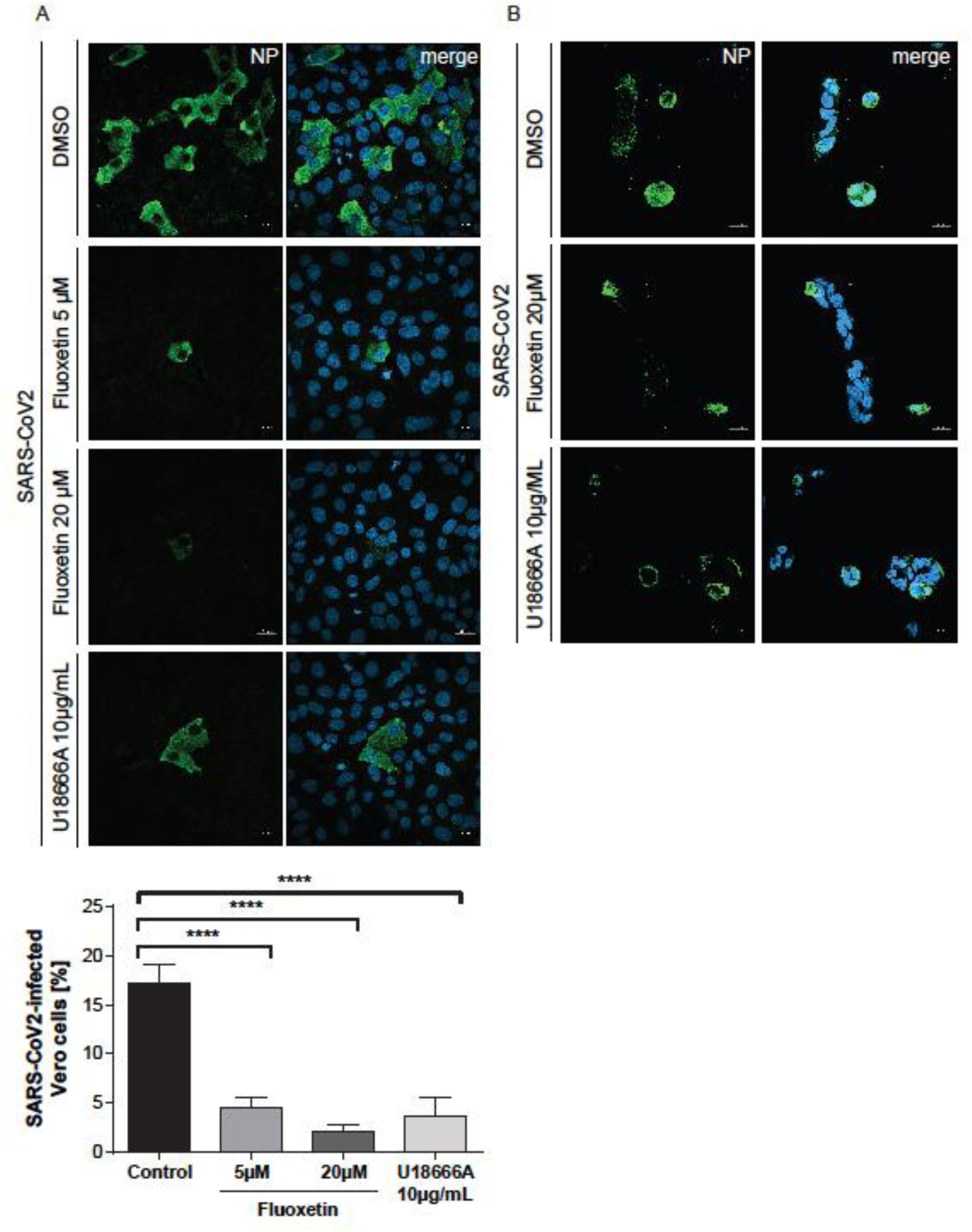
Impact of fluoxetine and U18666A on SARS-CoV-2 infection success within the first cycle of replication. (A) Vero and (B) Calu-3 cells preatreated with the drugs at the indicated concentrations were infected with SARS-CoV-2 at 1 MOI for 1 h. Nuclei were visualized with DAPI. To determine infection rates, NP-positive cells were detected by immunofluorescence imaging. Mean percentages ± SEM of NP-positive cells were calculated from 3 independent experiments. One-way ANOVA followed by by Dunnett’s multiple comparison test. ****p ≤ 0.0001.

## Discussion

In this study, we explored the use of small compounds that target the endolysosomal host-pathogen interface to interfere with SARS-CoV-2 infection. Enveloped viruses often exploit the host cell endocytic machinery for a safe transit into the cell. A critical step in the infection cycle is the fusion of the virus envelope with the cell membrane to to transfer the viral genome into the cytosol. Whereas IAV transfers the viral genome through fusion with the endosomal membrane [10,22–24], SARS-CoV-2 can fuse directly with the plasma membrane ^9^. However, recent evidence confirms that SARS-CoV-2 also uses endocytic uptake to enter the host cell ^25^ and full blocking of SARS-CoV-2 entry was only achieved when both the proteolytic activities of TMPRSS2 at the plasma membrane as well as of the endolysosomal cathepsins were blocked ^9^.

As endosomes move deeper into the cell, the pH in the lumen drops continuously, and this acidification distinguishes earlier from later stages of endosome maturation ^26,27^, thus indicating the successful progression of the virus-carrying endosomes into deeper parts of the cell. Therefore, endocytosed viruses often use the pH drop as a trigger for their endosomal escape ^10^. Moreover, the presentation of the viral fusogenic peptide is often supported through proteolytic cleavage by acid pH-dependent endolysosomal proteases ^10^. Proper endolysosomal function also requires the tight control of endolysosomal lipids, and both maintenance of the acidic pH and lipid balance are profoundly connected ^28,29^. Based on our earlier studies showing that the increase in endolysosomal endosomal cholesterol is embedded in the cell-autonomous interferon response against incoming influenza viruses, that this barrier has antiviral capacities when raised pharmacologically via repurposing of already licensed drugs, and is a druggable host cell target in a preclinical murine infection model ^12–14,19^, we investigated the impact of blocked endolysosomal cholesterol egress. Our results confirmed our previous results on IAV infection and show that increased endolysosomal cholesterol levels, together with the accompanying alterations in lumenal pH, also impair SARS-CoV-2 infection. Elevated cholesterol levels in the endolysosomal membranes render the vacuolar-type membrane ATPase (v-ATPase), the endolysosomal proton pump responsible for pH maintenance ^30^, inactive ^28^. Thus, the pH control within LELs might serve as a valuable antiviral target, a notion that is supported by the recent report on the inhibitory effect of the v-ATPase inhibitor bafilomycin A on SARS-CoV-2 cellular entry ^25^. However, the substantial toxicity prevents the clinical use of this macrolide antibiotic ^31^, and similar concerns limit the therapeutical use of U18666A ^32^. Dysbalanced endosomal cholesterol handling is also caused by inhibitory mutations in the endolysosomal enzyme acid sphingomyelinase (ASMase) which converts sphingomyelin to ceramide and phosphorylcholine in response to cell stress ^31^. ASMase activity is functionally inhibited by a large group of heterogeneous small compounds that are positively charged in an acidic environment such as the endolysosomal lumen. These so-called functional inhibitors of sphingomyelinase (FIASMAs) are clinically approved, generally well-tolerated, and widely used in human medicine for the treatment of a broad spectrum of pathological conditions ^16^. The FIASMA fluoxetine, trade-named Prozac, is a selective serotonin reuptake inhibitor that boomed in the 1980s and 1990s in the US and is commonly used to treat major depression and related disorders. Our results show that fluoxetine treatment was capable of inhibiting SARS-CoV-2 infection in a dose-dependent manner, with an EC_50_ value below 1 µM, and that the application of 10 µM fluoxetine severely reduced viral titers up to 99%. Fluoxetine-mediated pH neutralization was already seen at a low dose, whereas enhanced endolysosomal cholesterol pools were only visible when a higher dose was used.

Our results support the hypothesis that although there is quite some variation in the actual escape mechanisms ^11^, targeting the viral entry might serve as a target for antiviral therapy^33^. The intricate regulatory circuits that underly endolysosomal lipid balance and functionality are key elements functioning at the endolysosomal host-virus interface and are promising druggable targets for a wide variety of viruses and might be a fast and versatile approach to fight a broad range of pathogens with functionally similar modes of action. Because of the essential need for the host cell components, the infection cycle would have to be drastically altered to circumvent such host-directed therapeutics, and this approach is therefore considered much less likely to cause the development of resistance. The large variety of FIASMA pharmaceuticals offer a toolbox of potential antivirals for host-directed therapy, and exploring their use including their combination with drugs that directly target viral enzymes, might constitute a promising approach to repurpose these drugs as antivirals to counteract SARS-CoV-2 and COVID 19.

## Acknowledgements

We thank Jonathan Hentrey and Andreas Wilbers for help with the assays. This research was funded by grants from GERMAN RESEARCH FOUNDATION (DFG), CRC1009 “Breaking Barriers”, Project A06 (to U.R.) and B02 (to S.L.), CRC1348 “Dynamic Cellular Interfaces”, Project A11 (to U.R.), KFO342 TP6, Br5189/3-1 (to L.B.), Lu477/30-1 (to S.L.), INTERDISCIPLINARY CENTER FOR CLINICAL RESEARCH (IZKF) of the Münster Medical School, grant number Re2/022/20 and from the Innovative Medizinische Forschung (IMF) of the Münster Medical School, grant number SC121912 (to S.S.).

## Author contributions

Conceptualization, supervision and funding acquisition was done by U.R.; S.S. conceived the experiments and, together with J.G., A.M-Z., and N.K. carried out the experimental work and the data analysis. L.B., V.G., and S.L. provided resources and methodology. All authors have read and agreed to the submitted version of the manuscript.

## Figure Legends

**Suppl. Figure. 1.**
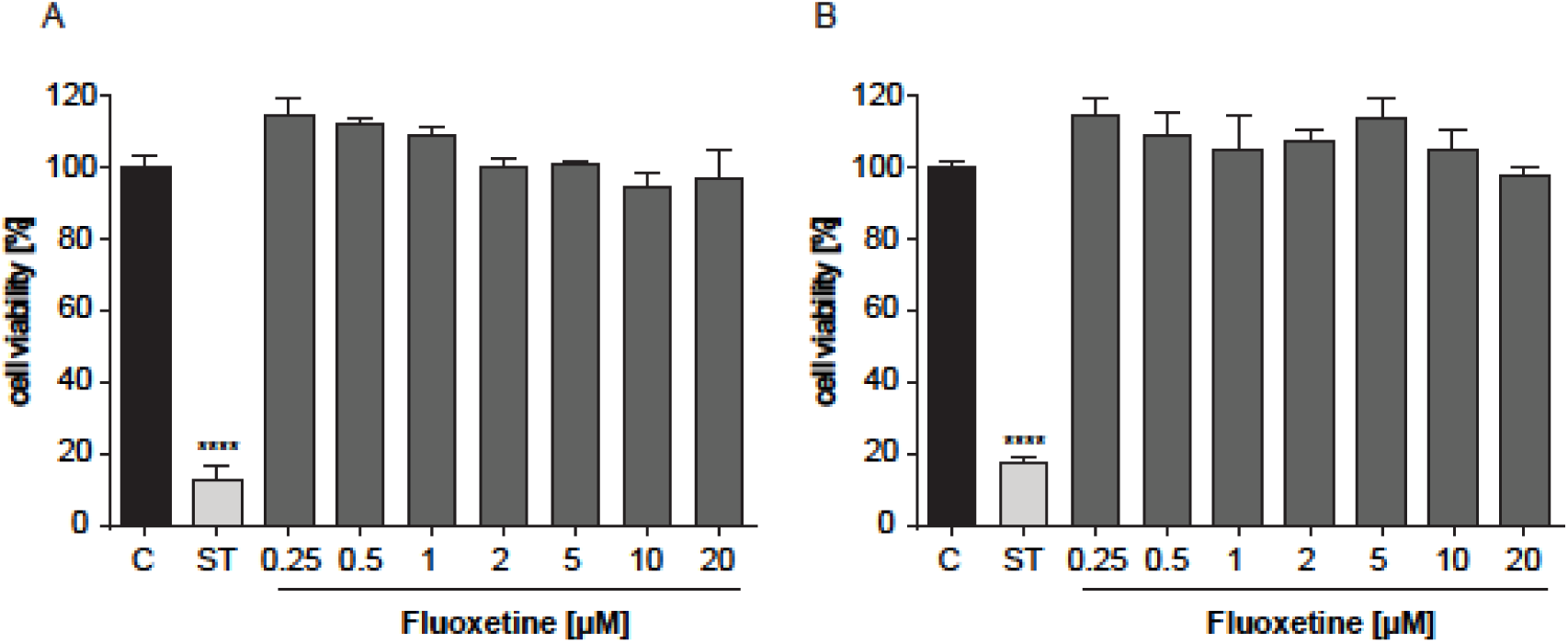
Analysis of cell viability. MTT assay of (A) Calu-3 and (B) vero cells treated with the solvent DMSO (C) or fluoxetine at the indicated concentrations for 48h. The protein kinase inhibitor staurosporine (ST), a strong inducer of cytotoxicity, served as a positive control. Data represent means ± SEM of three independent experiments; one-way ANOVA with Dunett’s multiple comparison tests, ****p ≤ 0.0001.

